## Supplemental Files for "Preparation of oxygen-sensitive proteins for high-resolution cryoEM structure determination using (an)aerobic blot-free vitrification"

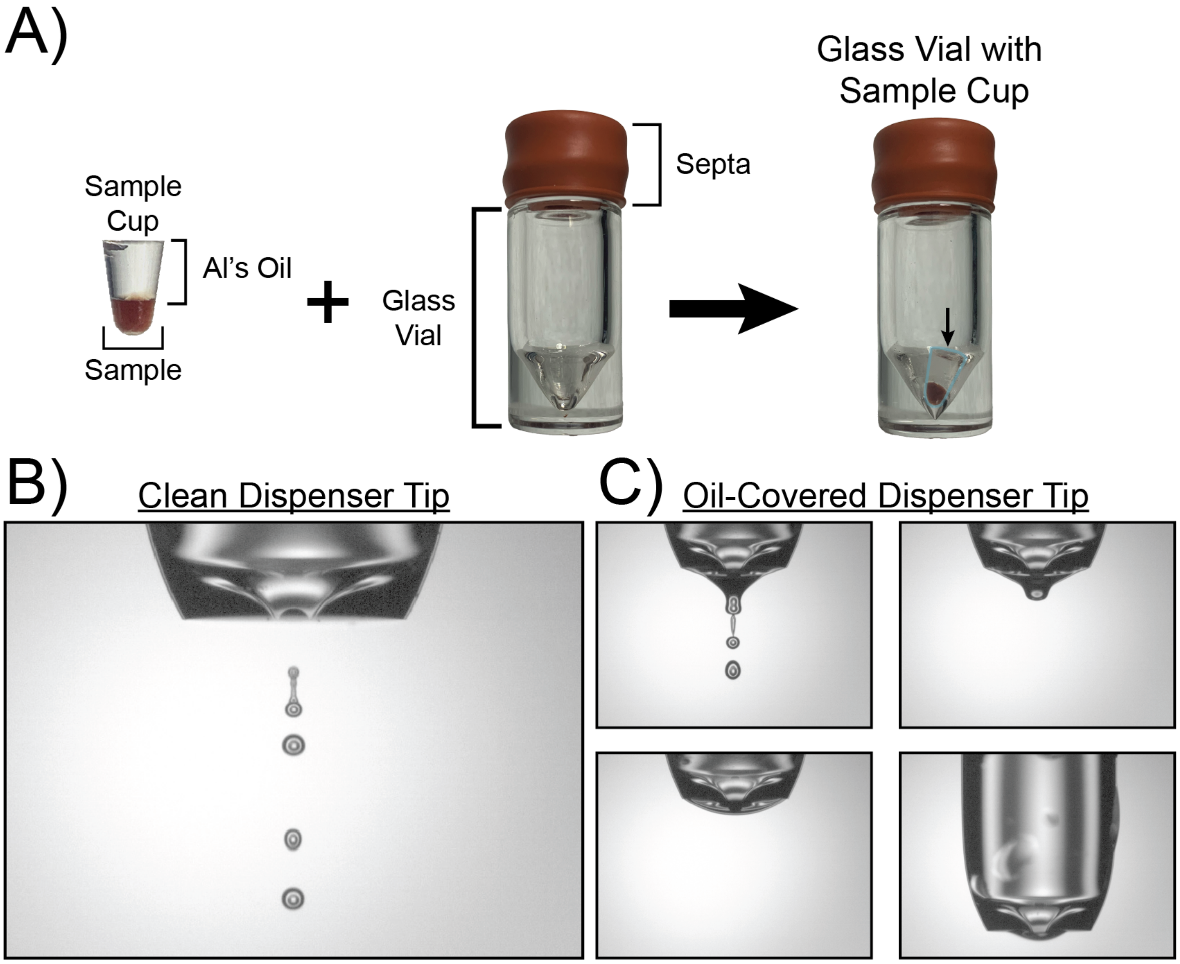


**Figure S1: Images of critical steps in the (an)aerobic workflow. A)** Images of a Hb sample in a chameleon sample cup with the protective Al's oil layer on top. The chameleon sample cup is prepared within an anaerobic environment and transferred into gas-tight glass vial and sealed with a rubber septum. **B-C** Images from the chameleon software displaying either a clean functioning dispenser tip **(B)** or a tip with residual oil on the outside **(C)** The residual oil can be removed using dispenser washing or wiping steps to yield a clean dispenser tip **(B)**.

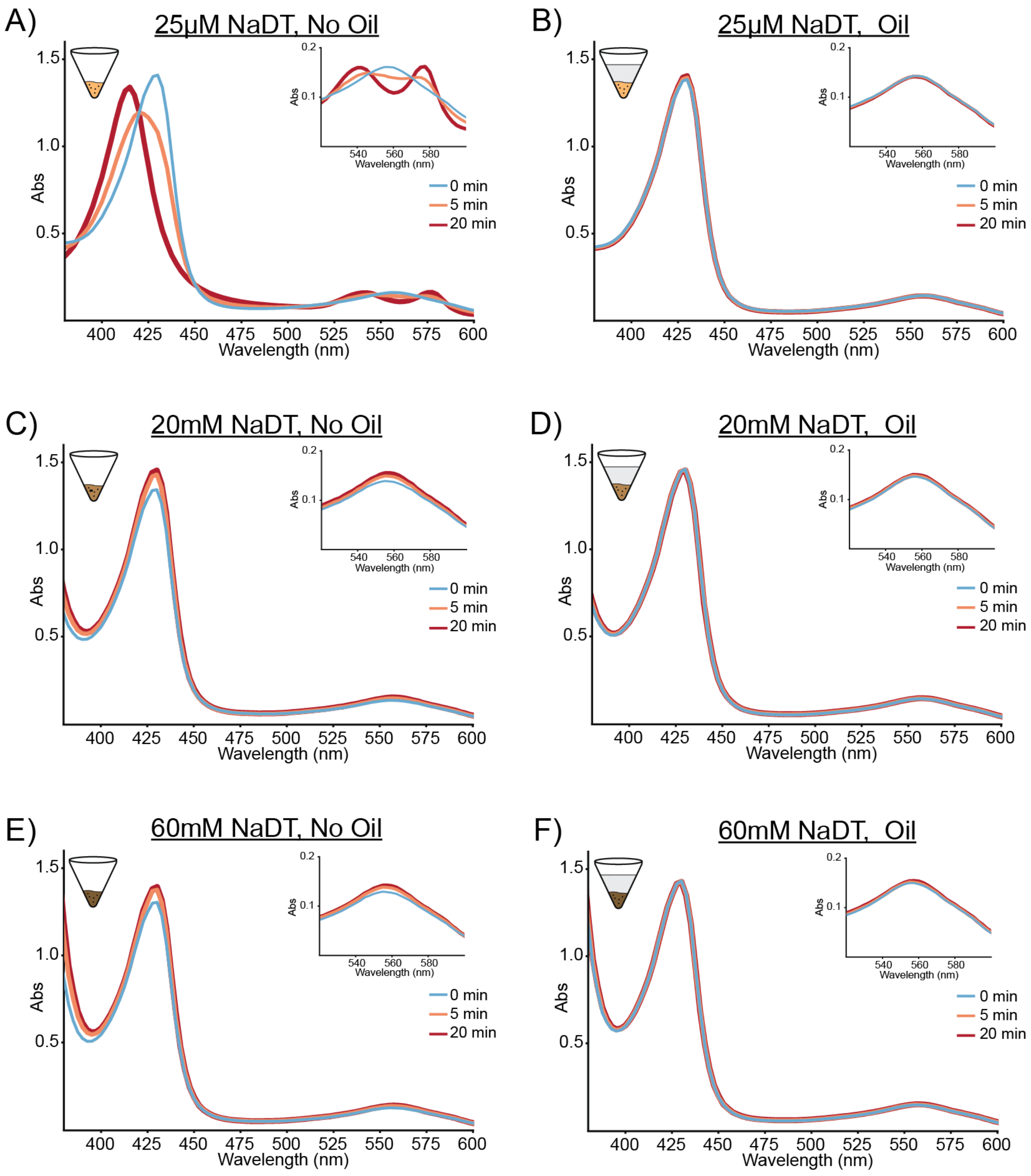


**Figure S2: UV-Vis-based oxygen perfusion assay using human deoxyHb under varying conditions.** For each sample, the anaerobic cuvette containing deoxyHb was opened to air and UV-Vis spectra were recorded from 350-600 nm over the course of 20 minutes. **A,B)** Timecourse of deoxyHb with 25 $\mu$M NaDT in sample buffer without and with a protective oil layer, respectively. **C,D)** Timecourse of deoxyHb with 20 mM NaDT in sample buffer without and with a protective oil layer, respectively. **E,F)** Timecourse of deoxyHb with 25 $\mu$M NaDT in sample buffer without and with a protective oil layer, respectively. For each assay, an inset of the 525-550 nm region is shown. For each assay, UV-Vis spectra at time 0 min (blue), 5 min (orange), and 20 min (red) are shown.

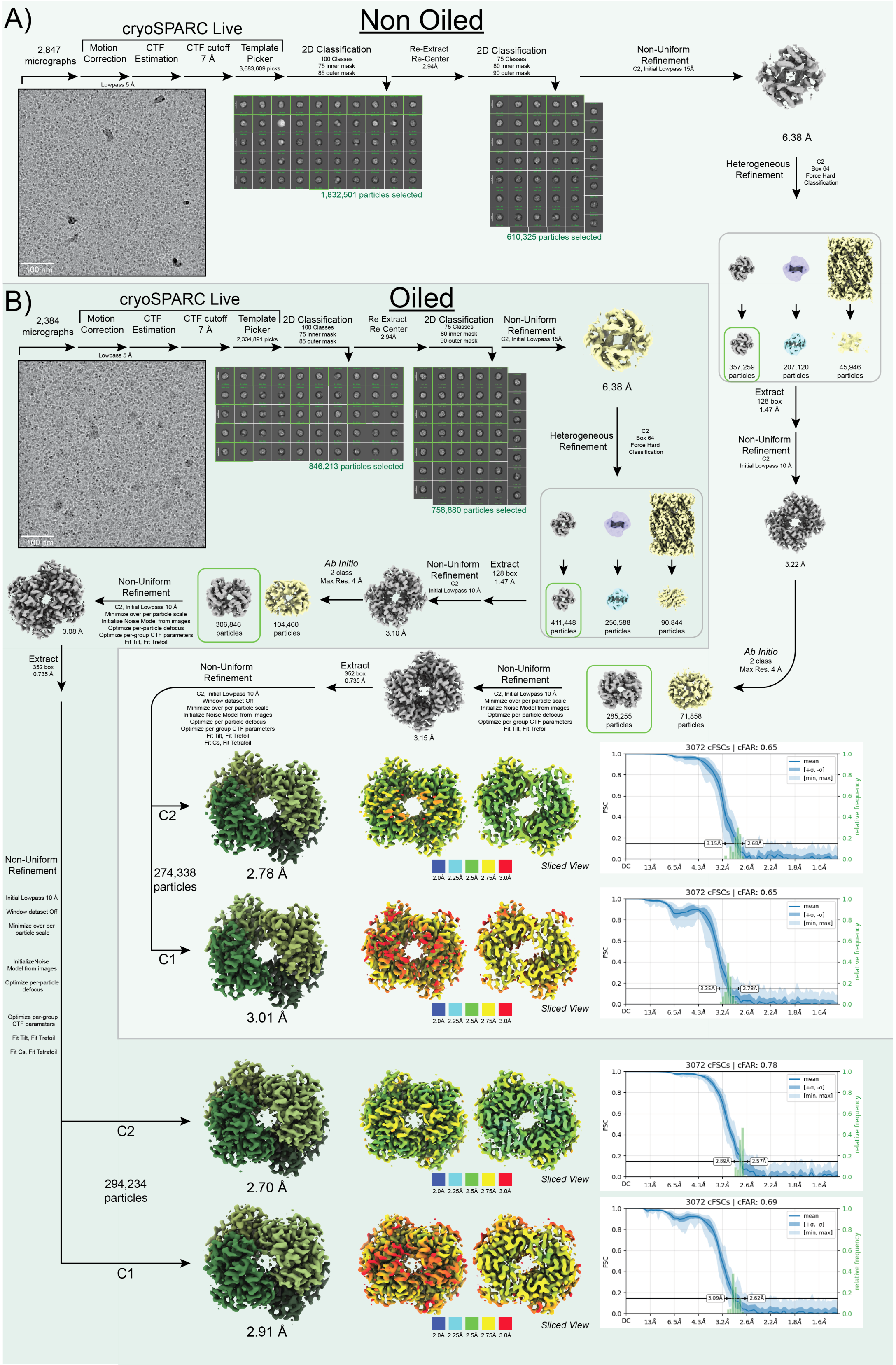

**Figure S3: CryoEM data processing workflow for non-oiled and oiled metHb structures. A)** Single-particle cryoEM data processing workflow for the 2.78 Å resolution non-oiled metHb structure. **B)** Single-particle cryoEM data processing workflow for the 2.71 Å resolution oiled metHb structure. 2,847 micrographs and 2,384 micrographs of non-oiled and oiled metHb, respectively, were collected and processed using similar strategies. Briefly, particle coordinates were obtained via template picking and extracted downsampled 4x prior to successive rounds of 2-D classification. Particles within the best classes were 3-D refined followed by a heterogeneous refinement using two Hb volumes and the 20S proteasome from *T. thermophilus* (EMDB-4877). Particles in the best class were carried downstream for 3-D refinement without downsampling with per-particle CTF and aberration refinements. For the final refinement, C1 and C2 symmetry were employed. Both non-oiled and oiled structures comprised similar final particle numbers, 275K versus 295K, respectively, and refined to similar FSC-estimated resolution with similar viewing distributions as estimated by 3-D FSC. Each of the final structures are colored by local resolution and the 3-D FSC plots are shown.

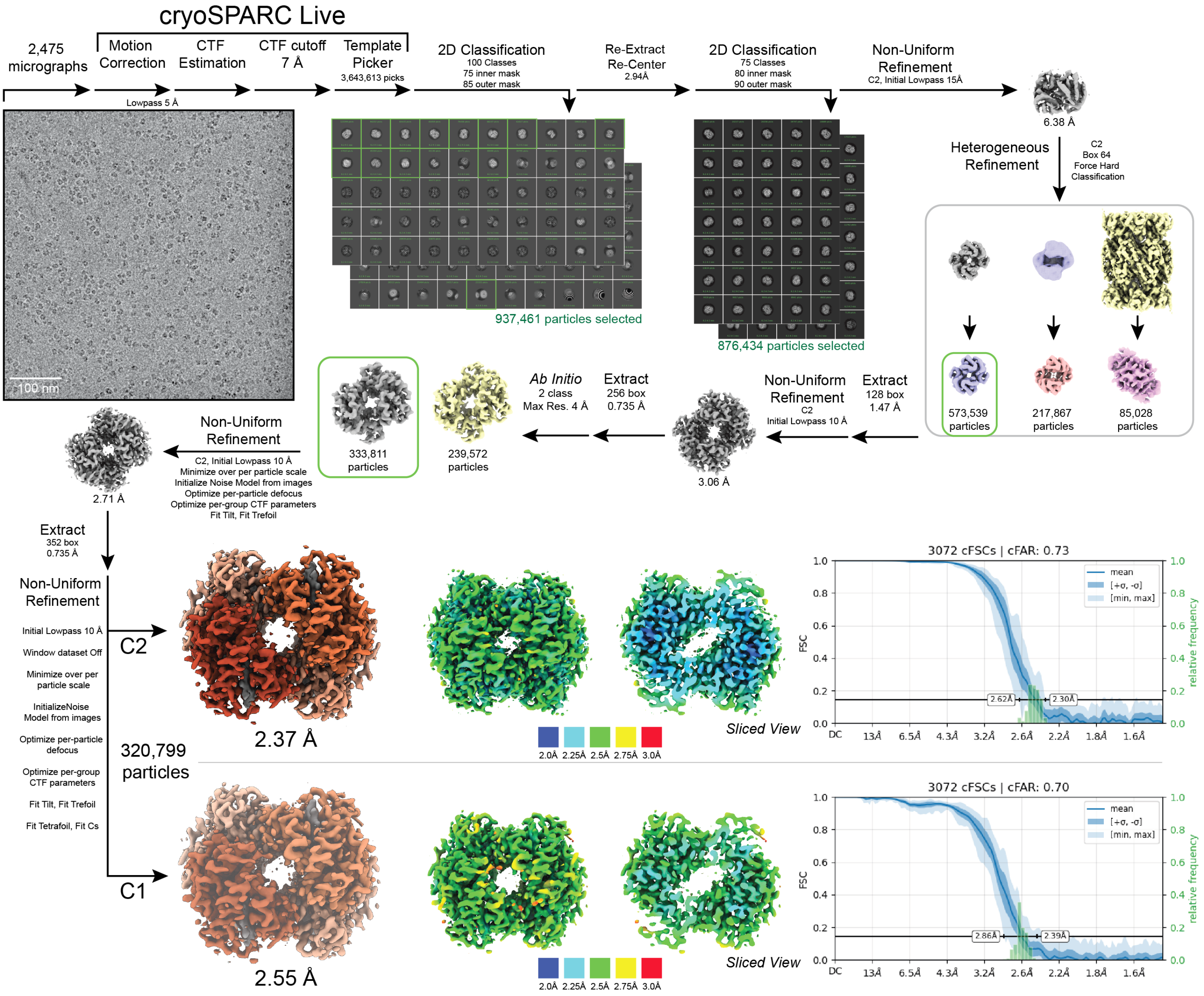


**Figure S4: CryoEM data processing workflow for the very low NaDT (25 $\mu$M), oxyHb structure.** Single-particle cryoEM data processing workflow for the 2.37 Å resolution oxyHb structure that was obtained in 25 $\mu$M NaDT under oil. 2,475 micrographs were collected and processed using a similar strategy as the metHb structures in **Fig S3**. Briefly, particle coordinates were obtained via template picking and extracted downsampled 4x prior to successive rounds of 2-D classification. Particles within the best classes were 3-D refined followed by a heterogeneous refinement using two Hb volumes and the 20S proteasome from *T. thermophilus* (EMDB-4877). Particles in the best class were carried downstream for 3-D refinement without downsampling with per-particle CTF and aberration refinements. For the final refinement, C1 and C2 symmetry were employed. Each of the final structures are colored by local resolution and the 3-D FSC plots are shown.

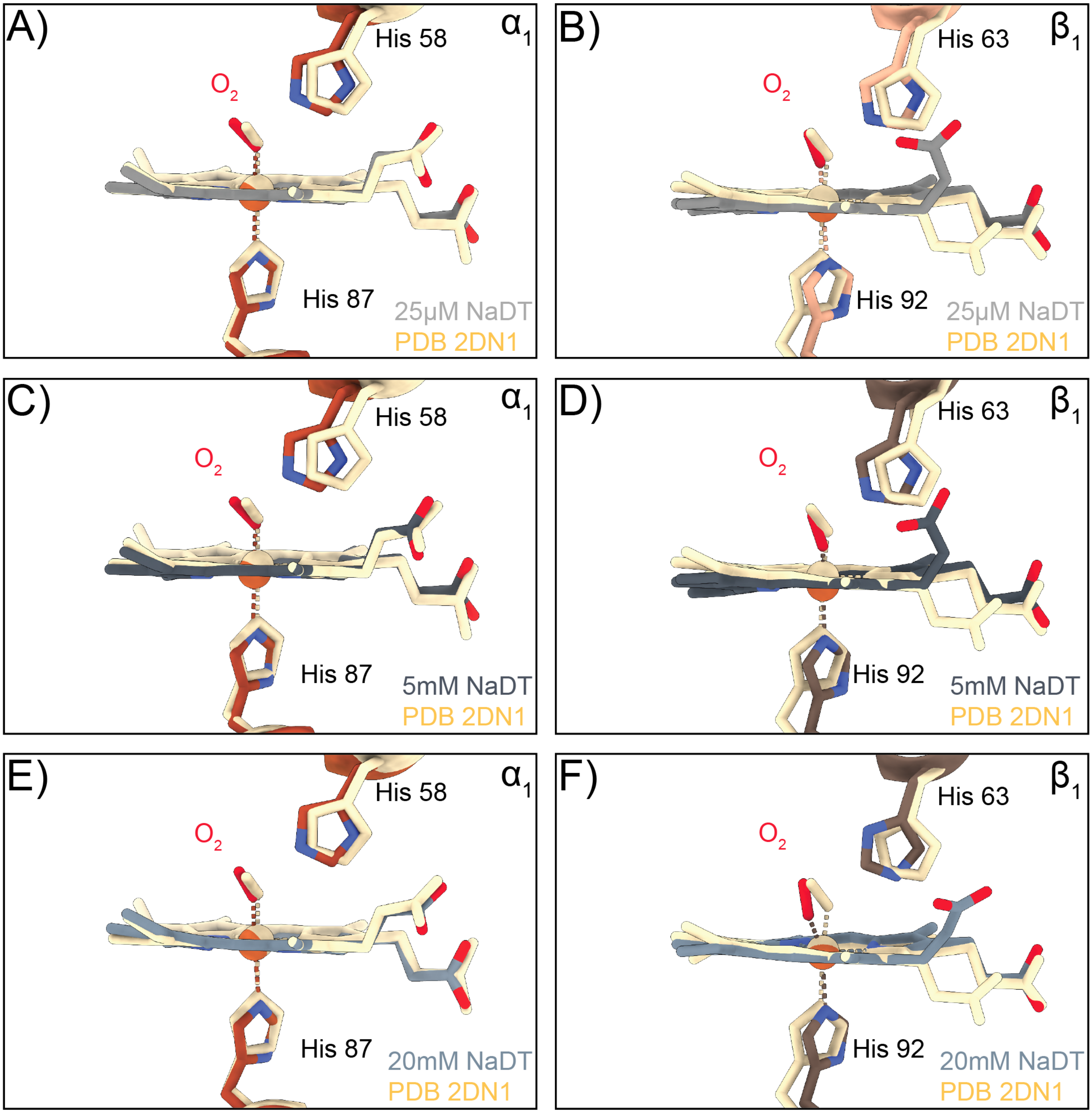


**Figure S5: Comparison of oxygen binding geometry in our oxyHb and partially-oxygenated Hb cryoEM structures vs oxyHb determined using X-ray crystallography structure (PDB: 2DN1). A-B)** The atomic models for the hemes of the $\alpha_1$ subunit **(A)** and $\beta_1$ subunit **(B)** (colored gray) and bound oxygen (colored red) overlapped with the heme and bound oxygen from the X-ray crystallography structure (PDB: 2DN1), shown in wheat detailing similar geometries **C-D)** The atomic models for the hemes of the $\alpha_1$ subunit **(C)** and $\beta_1$ subunit **(D)** (colored blue-gray) and bound oxygen (colored red) overlapped with the heme and bound oxygen from the X-ray crystallography structure (PDB: 2DN1), shown in wheat detailing similar geometries. The atomic models for $\alpha_1$H58, $\alpha_1$H87, $\beta_1$H63, and $\beta_1$H92 are also shown. **E-F)** The atomic models for the hemes of the $\alpha_1$ subunit **(E)** and $\beta_1$ subunit **(F)** (colored gray) and bound oxygen (colored wheat) overlapped with the heme and bound oxygen from the X-ray crystallography structure (PDB: 2DN1), shown in wheat detailing similar geometries. The atomic models for $\alpha_1$H58, $\alpha_1$H87, $\beta_1$H63, and $\beta_1$H92 are also shown.

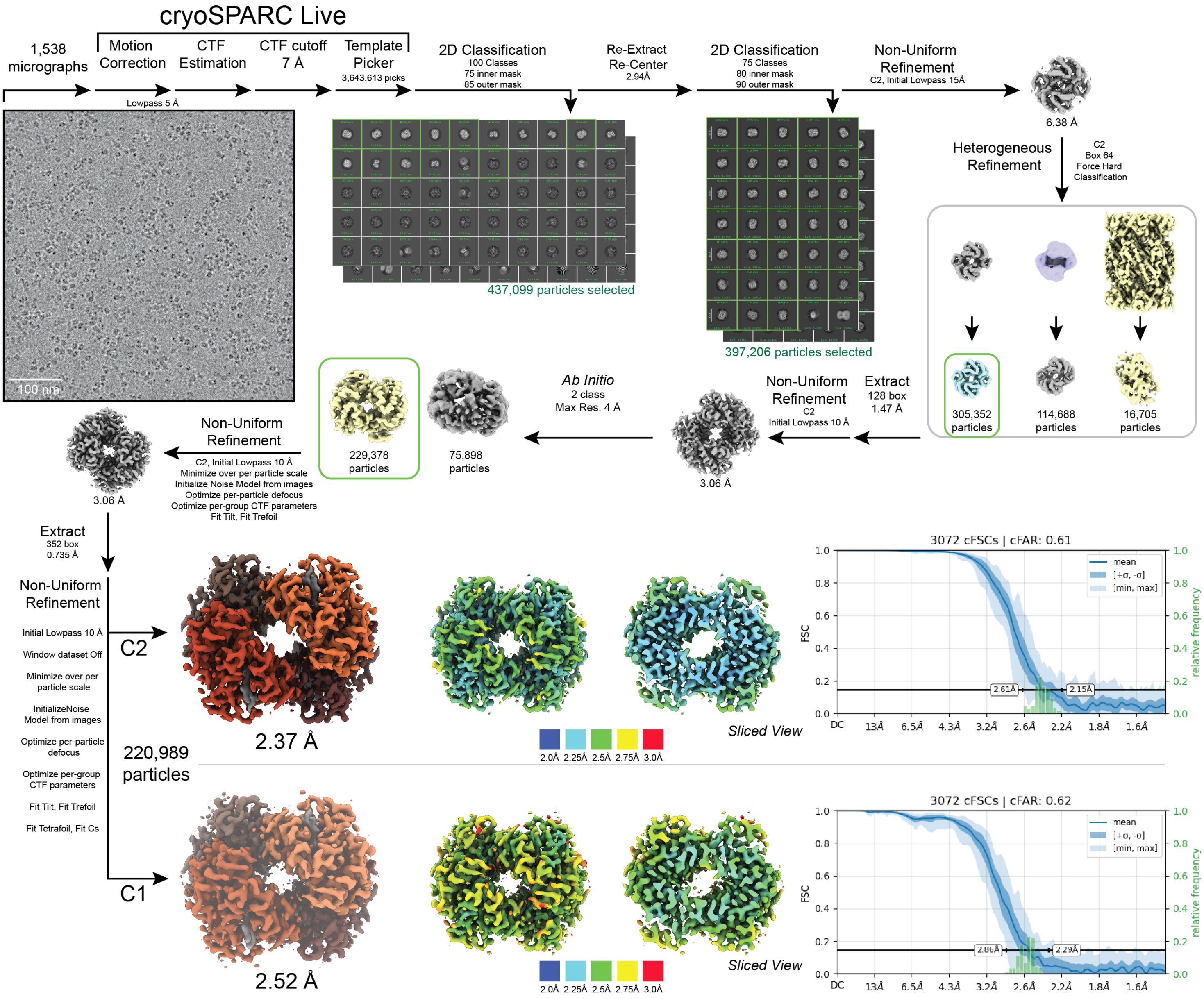


**Figure S6: CryoEM data processing workflow for the low NaDT (5 mM), oxyHb structure.** Single-particle cryoEM data processing workflow for the 2.37 Å resolution oxyHb structure that was obtained in 5 mM NaDT under oil. 1,538 micrographs were collected and processed using a similar strategy as the very low NaDT Hb structures in **Fig S4**. Briefly, particle coordinates were obtained via template picking and extracted downsampled 4x prior to successive rounds of 2-D classification. Particles within the best classes were 3-D refined followed by a heterogeneous refinement using two Hb volumes and the 20S proteasome from *T. thermophilus* (EMDB-4877). Particles in the best class were carried downstream for 3-D refinement without downsampling with per-particle CTF and aberration refinements. For the final refinement, C1 and C2 symmetry were employed. Each of the final structures are colored by local resolution and the 3-D FSC plots are shown.

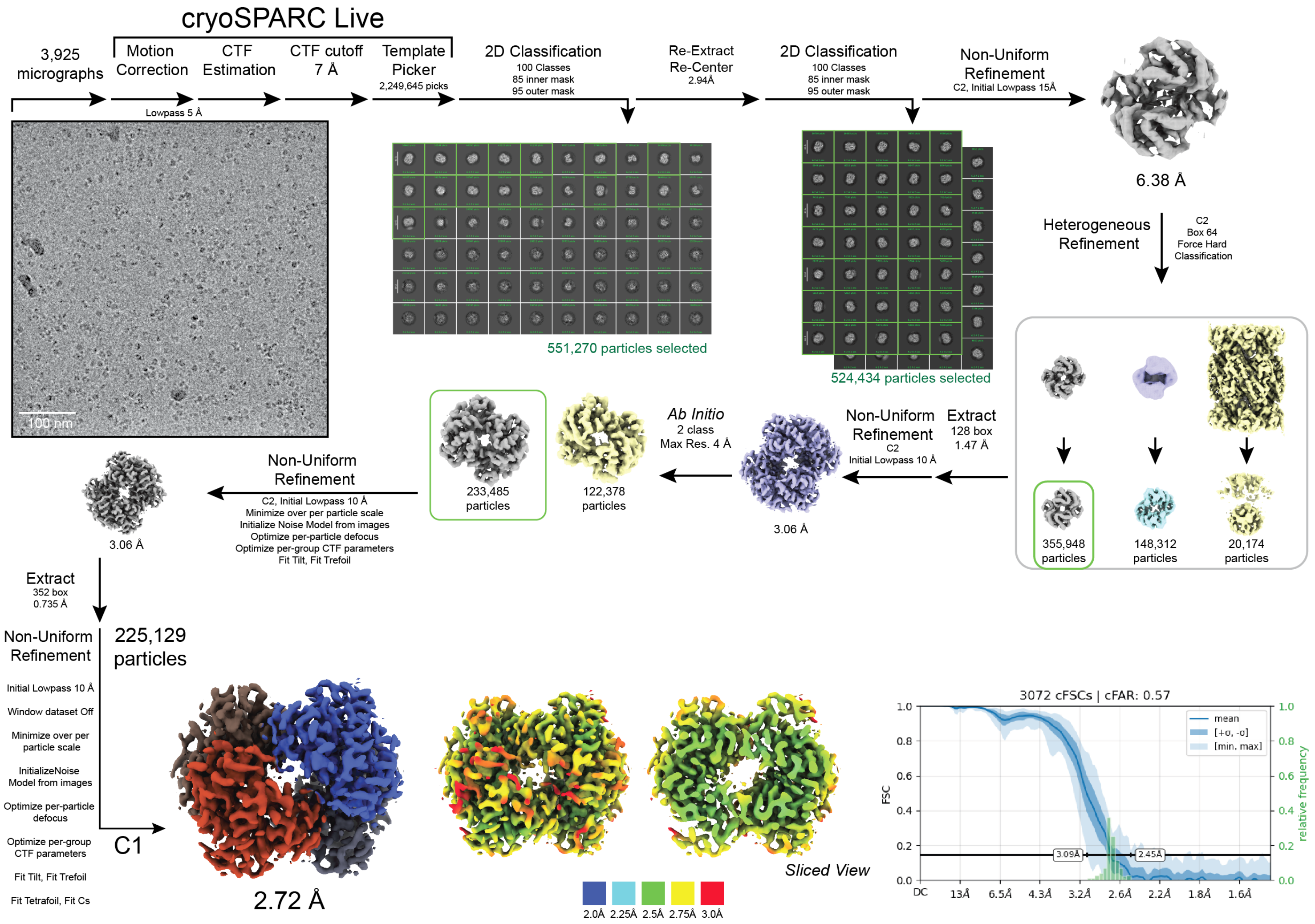


**Figure S7: CryoEM data processing workflow for the medium NaDT (20 mM), partially-oxygenated Hb structure.** Single-particle cryoEM data processing workflow for the 2.72 Å resolution partially-oxygenated (mixed) Hb structure that was obtained in 20 mM NaDT under oil. 3,925 micrographs were collected and processed using a similar strategy as the metHb structures in **Fig S3**. Briefly, particle coordinates were obtained via template picking and extracted downsampled 4x prior to successive rounds of 2-D classification. Particles within the best classes were 3-D refined followed by a heterogeneous refinement using two Hb volumes and the 20S proteasome from *T. thermophilus* (EMDB-4877). Particles in the best class were carried downstream for 3-D refinement without downsampling with per-particle CTF and aberration refinements. For the final refinement, C1 symmetry was employed. The final structure is colored by local resolution and the 3-D FSC plot is shown.

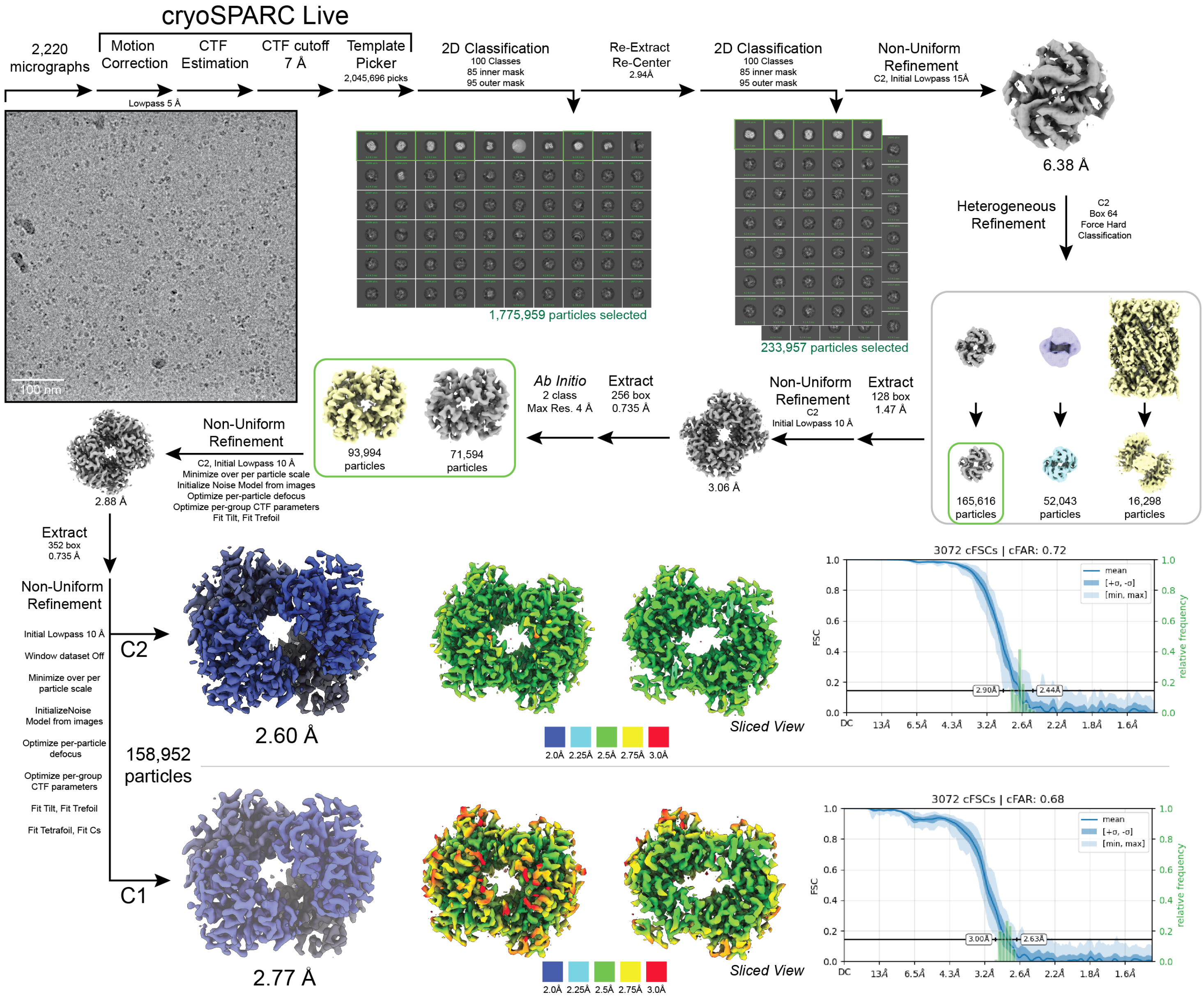


**Figure S8: CryoEM data processing workflow for the high NaDT (60 mM), deoxyHb structure.** Single-particle cryoEM data processing workflow for the 2.60 Å resolution deoxyHb structure that was obtained in 60 mM NaDT under oil. 2,220 micrographs were collected and processed using a similar strategy as the metHb structures in **Fig S3**. Briefly, particle coordinates were obtained via template picking and extracted downsampled 4x prior to successive rounds of 2-D classification. Particles within the best classes were 3-D refined followed by a heterogeneous refinement using two Hb volumes and the 20S proteasome from *T. thermophilus* (EMDB-4877). Particles in the best class were carried downstream for 3-D refinement without downsampling with per-particle CTF and aberration refinements. For the final refinements, C1 and C2 symmetry were employed. Each of the final structures are colored by local resolution and the 3-D FSC plots are shown.

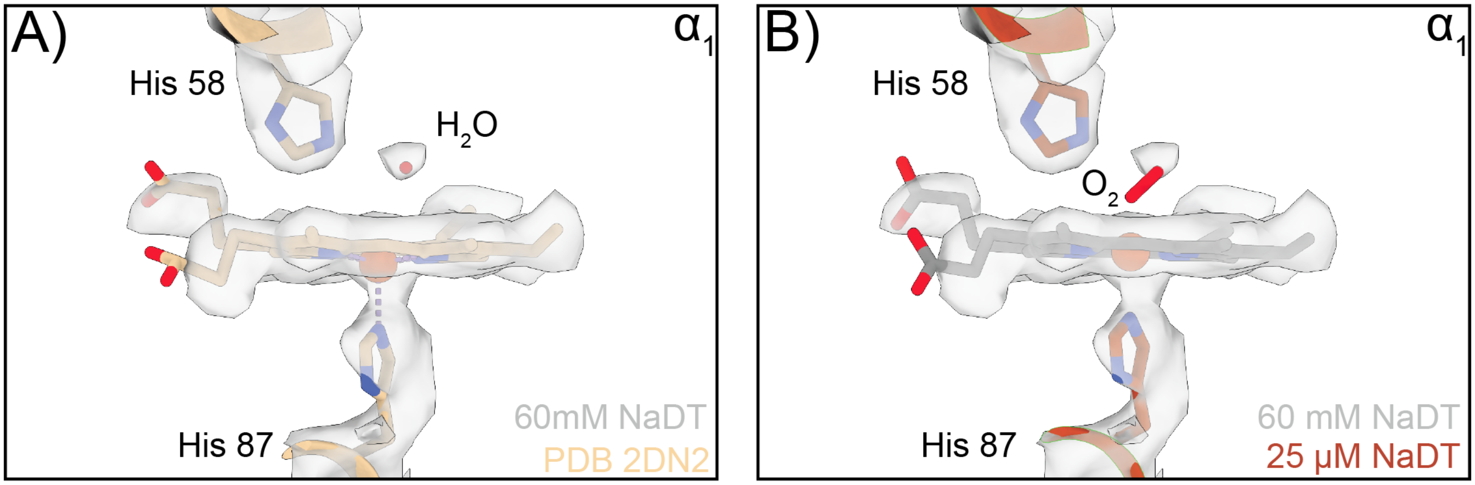


**Figure S9: Water density in the distal pocket of the deoxyHb structure. A)** CryoEM density of the 60 mM NaDT deoxyHb structure with the 1.25 Å deoxyHb X-ray atomic model (PDB: 2DN2) docked in place showing a well-defined water molecule above the ferrous unliganded heme of the $\alpha_1$ subunit. **B)** Same cryoEM density from **(A)** with our oxyHb atomic model docked in showing the poor fit of the oxygen ligand. The atomic models for $\alpha_1$H58, $\alpha_1$H87 are also shown.

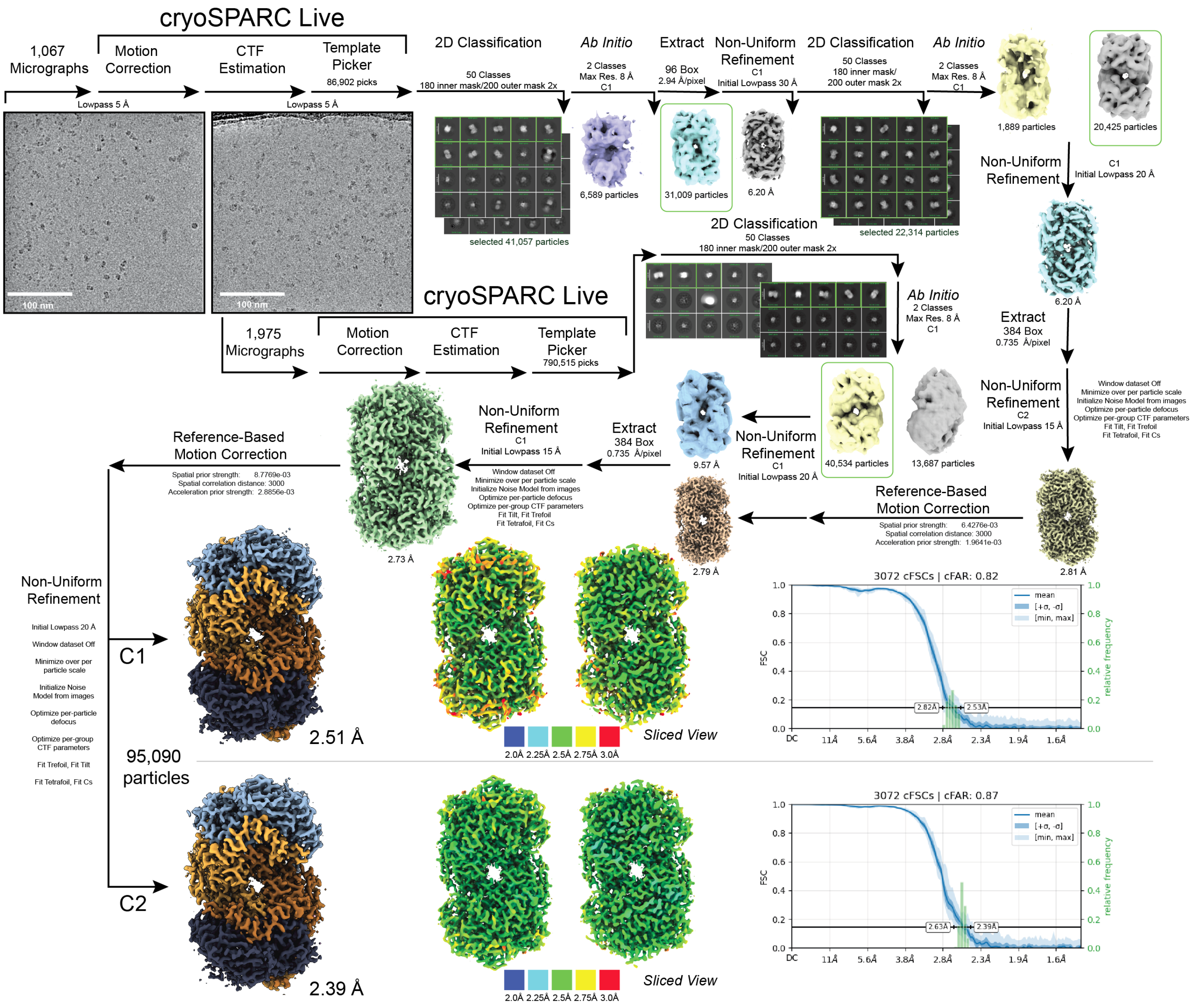


**Figure S10: CryoEM data processing workflow for the oxidized ($P^{2+}$) *Av*MoFeP structure.** Single-particle cryoEM data processing workflow for the 2.39 Å resolution oxidized ($P^{2+}$) *Av*MoFeP structure that was obtained in without the protective oil layer and without NaDT in the cryoEM buffer. 1,067 and 1,975 micrographs were collected across two collections and processed using a similar strategies before being combined at the end. Briefly, particle coordinates were obtained via template picking and extracted downsampled 4x prior to successive rounds of 2-D classification. Particles within the best classes were subjected to a 2-class *ab initio* model generation. Particles in the best class were then subjected to another round of 2-D classification and 2-class *ab initio* prior to 3-D refinement of the best particles without downsampling. Reference-based motion correction was performed for each particle set before combining for a non-uniform refinement with per-particle CTF and aberration refinements. For the final refinements, C1 and C2 symmetry were employed. Each of the final structures are colored by local resolution and the 3-D FSC plots are shown.

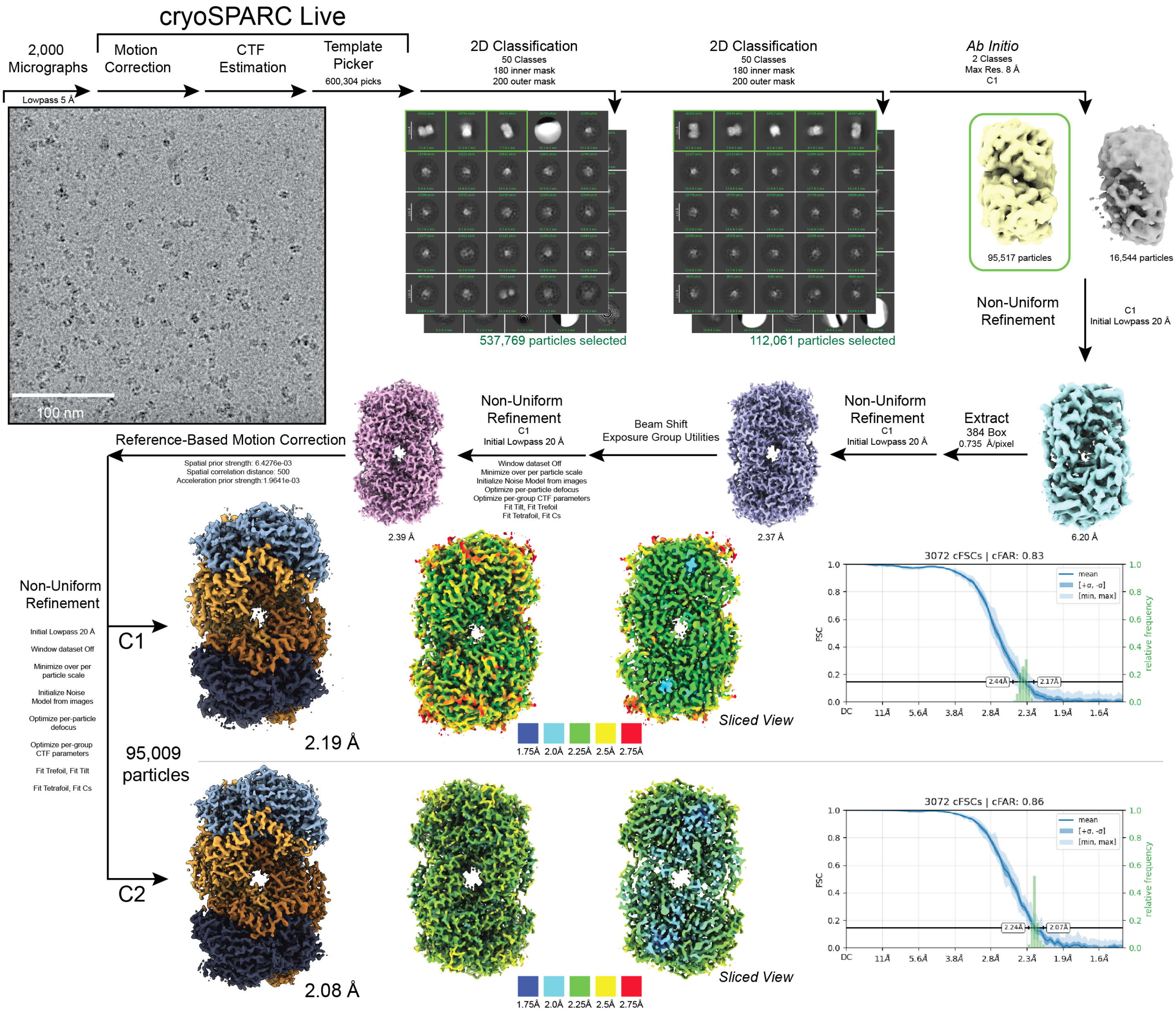


**Figure S11: CryoEM data processing workflow for the reduced ($P^N$) *Av*MoFeP structure.** Single-particle cryoEM data processing workflow for the 2.08 Å resolution reduced ($P^N$) *Av*MoFeP structure that was obtained with the protective oil layer and 20 mM NaDT in the cryoEM buffer. 2,000 micrographs were collected processed using a similar strategy as the oxidized ($P^{2+}$) MoFeP structure. For the final refinements, C1 and C2 symmetry were employed. Each of the final structures are colored by local resolution and the 3-D FSC plots are shown.

| | No Oil Hb | | Oil Hb | | "No" NaDT | | Low NaDT | | Partial Oxy Hb | | DeOxy Hb | |
|---|---|---|---|---|---|---|---|---|---|---|---|---|
| **Data Collection** | | | | | | | | | | | | |
| Magnification | 165 kx | | 165 kx | | 165 kx | | 165 kx | | 165 kx | | 165 kx | |
| Voltage (kV) | 300 | | 300 | | 300 | | 300 | | 300 | | 300 | |
| Spherical Aberration (mm) | 2.7 | | 2.7 | | 2.7 | | 2.7 | | 2.7 | | 2.7 | |
| Electron Exposure ($e^-/\text{Å}^2$) | 60 | | 60 | | 60 | | 60 | | 60 | | 60 | |
| Defocus range ($\mu$m) | -1.0 to -2.5 | | -1.0 to -2.5 | | -1.0 to -2.5 | | -1.0 to -2.5 | | -1.0 to -2.5 | | -1.0 to -2.5 | |
| Pixel size (Å, Physical/Digital) | 0.735 | | 0.735 | | 0.735 | | 0.735 | | 0.735 | | 0.735 | |
| Energy Filter Slit Width (eV) | 10 | | 10 | | 10 | | 10 | | 10 | | 10 | |
| **Map Statistics and Post-Processing** | | | | | | | | | | | | |
| Symmetry imposed | C1 | C2 | C1 | C2 | C1 | C2 | C1 | C2 | C1 | C2 | C1 | C2 |
| Map Resolution (Å) | 3.01 | 2.78 | 2.91 | 2.7 | 2.55 | 2.37 | 2.52 | 2.37 | 2.72 | 2.56 | 2.75 | 2.61 |
| Local resolution range for 75% of voxels | 7.069 | 6.294 | 6.908 | 5.975 | 6.521 | 5.798 | 6.769 | 5.901 | 6.983 | 5.971 | 7.095 | 6.344 |
| Local resolution range (model) | 2.6 - 41.1 | 2.4 - 26.9 | 2.5 - 42.7 | 2.3 - 22.6 | 1.6 - 38.8 | 2.1 - 27.6 | 2.2 - 39.0 | 2.1 - 34.5 | 2.4 - 42.7 | 2.2 - 38.4 | 1.6 - 41.6 | 2.3 - 25.4 |
| Map sharpening B factor ($\text{Å}^2$) | 118.3 | 111.9 | 112.3 | 110.2 | 89.2 | 85.9 | 82.7 | 80.5 | 91.9 | 91.1 | 89.5 | 93.4 |
| Map sharpening method | Resolve | Resolve | Resolve | Resolve | Resolve | Resolve | Resolve | Resolve | Resolve | Resolve | Resolve | Resolve |
| Q-Score | 0.7 | 0.74 | 0.73 | 0.76 | 0.78 | 0.81 | 0.78 | 0.78 | 0.75 | 0.78 | 0.76 | 0.78 |
| Movies | 2847 | 2847 | 2384 | 2384 | 2475 | 2475 | 1538 | 1538 | 3925 | 3925 | 2220 | 2220 |
| **Model Statistics and Validation** | | | | | | | | | | | | |
| Model composition | | | | | | | | | | | | |
| Non-hydrogen atoms | 4543 | 4577 | 4547 | 4584 | 4604 | 4591 | 4853 | 4950 | 4587 | 4862 | 4580 | 4601 |
| Protein | 568 | 568 | 568 | 568 | 568 | 568 | 568 | 568 | 568 | 568 | 568 | 568 |
| Nucleic acids | 0 | 0 | 0 | 0 | 0 | 0 | 0 | 0 | 0 | 0 | 0 | 0 |
| Ligand1 | HEM: 4 | HEM: 4 | HEM: 4 | HEM: 4 | HEM: 4 | HEM: 4 | HEM: 4 | HEM: 4 | HEM: 4 | HEM: 4 | HEM: 4 | HEM: 4 |
| Waters | 51 | 85 | 55 | 92 | 104 | 91 | 353 | 450 | 91 | 366 | 88 | 109 |
| R.M.S deviations | | | | | | | | | | | | |
| Length (Å) | 0.004 | 0.039 | 0.033 | 0.005 | 0.047 | 0.022 | 0.003 | 0.003 | 0.024 | 0.003 | 0.024 | 0.039 |
| Angles (°) | 0.602 | 1.496 | 1.215 | 0.934 | 2.128 | 1.218 | 0.512 | 0.531 | 1.331 | 0.673 | 1.466 | 1.940 |
| MolProbity score | 1.24 | 1.27 | 1.30 | 1.36 | 1.30 | 1.18 | 1.36 | 1.48 | 1.23 | 1.27 | 1.18 | 1.19 |
| MolProbity Clashscore | 4.72 | 5.06 | 5.61 | 4.60 | 5.50 | 3.93 | 6.06 | 5.38 | 4.49 | 5.16 | 3.93 | 4.04 |
| CaBLAM (% outliers) | 0.91 | 0.93 | 0.73 | 0.72 | 0.75 | 0.73 | 0.54 | 0.72 | 0.36 | 0.72 | 0.55 | 0.74 |
| Rotamer outliers (%) | 0.66 | 0.44 | 0.44 | 1.32 | 0.44 | 0.88 | 1.10 | 1.76 | 0.44 | 0.00 | 0.44 | 0.00 |
| Cis peptides (#, %) | 0.0/0.0 | 0.0/0.0 | 0.0/0.0 | 0.0/0.0 | 0.0/0.0 | 0.0/0.0 | 0.0/0.0 | 0.0/0.0 | 0.0/0.0 | 0.0/0.0 | 0.0/0.0 | 0.0/0.0 |
| Ramachandran Plot | | | | | | | | | | | | |
| Favored | 98.92 | 98.18 | 98.74 | 97.86 | 98.72 | 99.82 | 99.29 | 99.64 | 99.28 | 99.11 | 99.1 | 99.46 |
| Allowed | 1.08 | 1.82 | 1.26 | 2.14 | 1.28 | 0.18 | 0.71 | 0.36 | 0.72 | 0.89 | 0.9 | 0.54 |
| Outliers | 0.0 | 0.0 | 0.0 | 0.0 | 0.0 | 0.0 | 0.0 | 0.0 | 0.0 | 0.0 | 0.0 | 0.0 |

**Table 1:** CryoEM data collection and refinement statistics of No Oil, Oil, Very Low NaDT, Low NaDT, Partial Oxy, and DeOxy Hb.

| | Oxidized MoFeP | | Reduced MoFeP | |
|---|---|---|---|---|
| **Data Collection** | | | | |
| Magnification | 165 kx | | 165 kx | |
| Voltage (kV) | 300 | | 300 | |
| Spherical Aberration (mm) | 2.7 | | 2.7 | |
| Electron Exposure ($e^-/Å^2$) | 60 | | 60 | |
| Defocus range ($\mu$m) | -0.75 to -2.5 | | -0.75 to -2.5 | |
| Pixel size (Å, Physical/Digital) | 0.735 | | 0.735 | |
| Energy Filter Slit Width (eV) | 10 | | 10 | |
| **Map Statistics and Post-Processing** | | | | |
| Symmetry imposed | C1 | C2 | C1 | C2 |
| Map Resolution (Å) | 2.51 | 2.39 | 2.19 | 2.08 |
| Local resolution range for 75% of voxels | 6.69 | 6.034 | 6.273 | 5.658 |
| Local resolution range (model) | 2.1 - 39.0 | 1.6 - 38.1 | 1.9 - 35.2 | 1.7 - 35.2 |
| Map sharpening B factor ($Å^2$) | 65.2 | 68.3 | 51.0 | 51.8 |
| Map sharpening method | Resolve | Resolve | Resolve | Resolve |
| Q-Score | 0.8 | 0.82 | 0.85 | 0.85 |
| Movies | 2002 | 2002 | 2000 | 2000 |
| **Model Statistics and Validation** | | | | |
| Model composition | | | | |
| Non-hydrogen atoms | 17271 | 17365 | 17462 | 17763 |
| Protein | 1998 | 1998 | 1998 | 1998 |
| Nucleic acids | 0 | 0 | 0 | 0 |
| Ligand1 | ICS: 2 | ICS: 2 | ICS: 2 | ICS: 2 |
| Ligand2 | CLF: 2 | CLF: 2 | CLF: 2 | CLF: 2 |
| Ligand3 | HCA: 2 | HCA: 2 | HCA: 2 | HCA: 2 |
| Ligand4 | FE: 2 | FE: 2 | FE: 2 | FE: 2 |
| Waters | 1244 | 1338 | 1435 | 1736 |
| R.M.S deviations | | | | |
| Length (Å) | 0.003 | 0.004 | 0.002 | 0.003 |
| Angles (°) | 0.509 | 0.586 | 0.497 | 0.560 |
| MolProbity score | 1.66 | 1.59 | 1.40 | 1.82 |
| MolProbity Clashscore | 6.85 | 9.63 | 6.41 | 14.96 |
| CaBLAM (% outliers) | 0.61 | 0.76 | 0.55 | 0.71 |
| Rotamer outliers (%) | 1.57 | 1.04 | 1.04 | 0.81 |
| Cis peptides (#, %) | 6.5/0.1 | 6.5/0.1 | 6.5/0.1 | 6.5/0.1 |
| Ramachandran Plot | | | | |
| Favored | 97.29 | 97.69 | 97.84 | 97.24 |
| Allowed | 2.71 | 2.31 | 2.16 | 2.76 |
| Outliers | 0.0 | 0.0 | 0.0 | 0.0 |

**Table 2:** CryoEM data collection and refinement statistics of Oxidized and Reduced MoFeP
